# Translationally relevant transcriptomic alterations in mouse ischemic cerebral microvessels

**DOI:** 10.1101/829820

**Authors:** Keri Callegari, Sabyasachi Dash, Hiroki Uchida, Yunkyoung Lee, Akira Ito, Tuo Zhang, Jenny Xiang, Teresa Sanchez

## Abstract

Increasing evidence implicates cerebral microvascular dysfunction in the pathophysiology of numerous central nervous system pathologies, including stroke. Understanding the molecular alterations in cerebral microvessels in these conditions will provide original opportunities for scientific investigation at the pre-clinical and clinical levels. In this study, we conducted a novel genome-wide transcriptomic analysis of microvessels in a mouse model of transient focal cerebral ischemia. Using a publicly available human ischemic stroke dataset, we identified shared alterations in our microvessel dataset with implications for human pathophysiology. From this unbiased analysis, we report predicted alterations in inter- and intra-cellular signaling, emphasizing perturbations in genes involved in blood brain barrier function, endothelial cell activation and metabolism. Furthermore, our study unveiled previously unreported gene expression changes associated with altered sphingolipid metabolism. Altogether, our results have identified microvessel-specific transcriptomic changes in a number of translationally relevant pathways that support the targeting of these pathways in preclinical studies. The data shared here provide a resource for future investigation of translationally relevant pathways in ischemic stroke.

## INTRODUCTION

Despite many decades of research, stroke is still a leading cause of mortality and disability worldwide[1, 2]. While stroke research has focused largely on development of neuroprotective agents, none of these drugs unequivocally showed an improvement in clinical outcomes[3, 4]. Thus, there is a need to develop novel therapeutic strategies[5]. There is increasing evidence that cerebral microvascular dysfunction plays a critical role in the exacerbation of neurovascular injury in stroke[6–14]. The cerebrovascular endothelium, in coordination with pericytes [15, 16] and astrocytes[17] plays a critical role in the maintenance of the blood brain barrier (BBB). As the primary barrier between systemic blood supply and the central nervous system, it has great therapeutic potential [18] [19, 20].

Stroke induced alterations in BBB integrity and function are important contributors to brain injury related to hypoxia and neuroinflammation [21]. In the acute phase of stroke, pro-inflammatory cytokines (TNF-α, IL-1β) are released and trigger the induction of matrix metalloproteinases 3 and 9 (MMP-3/MMP-9) that degrade the basal lamina and contribute to endothelial activation and BBB permeability[22]. BBB dysfunction exacerbates neurovascular ischemic injury by allowing the entrance of neurotoxic plasma components into the brain parenchyma, increasing intra-cerebral pressure with the risk of brain herniation and vessel compression, further compromising blood flow to the brain [6, 7, 10, 23, 24]. Endothelial activation is marked by dysregulated endothelial function culminating in microvascular thrombosis and tissue damage with progressive cell death. The pathways leading to and resulting from endothelial dysfunction in ischemia require extensive investigation to broaden our understanding of the molecular mechanisms governing disease progression. By expanding our comprehension of the underlying biology at the neurovascular level, we can develop potential therapeutics for clinical intervention. There is an increasingly dire need for novel targeted approaches to prevent BBB disruption in the acute and long-term stroke-related pathophysiology.

In order to expand our understanding of how cerebral microvascular dysfunction contributes to ischemic pathology, in the present study, we conducted an unbiased experiment to determine transcriptomic changes in cerebral microvessels after stroke that are relevant to human stroke pathology. We elucidated several signaling pathways relevant to inflammation and metabolic stress in the BBB and identified targetable pathways enriched in microvessel preparations. From this unbiased analysis, we report predicted alterations in genes involved in blood brain barrier dysfunction, endothelial cell activation and metabolism, which are sustained in human stroke lesions highlighting the contribution of these pathways to chronic pathophysiology. In addition, given the encapsulating nature of our RNA-sequencing data, we explored transcriptomic changes relevant to sphingolipid metabolism to uncover novel mechanisms for sphingosine-1-phosphate signaling alterations in stroke. Our data provide support for future preclinical studies to explore neurovascular therapies pertinent to human disease.

## RESULTS

### Transcriptomic changes in cerebral microvessels after tMCAO

In order to examine gene expression changes at the neurovascular level, microvessels were isolated from the cerebral cortices of mice subjected to transient middle cerebral artery occlusion (tMCAO) or sham surgery (Figure 1A). This protocol was completed as previously described [25]. RNA was isolated from these microvessels and sequenced on an Illumina platform (n=4). Outliers were excluded similarly across all samples and tMCAO samples clustered with similar gene expression patterns (Supplemental Figure 1). Interestingly, a mild subgrouping within sham microvessels was observed (Supplemental Figure 1C-D). General examination of transcript expression alterations revealed 18,491 out of 32,129 genes were altered, of which 6,291 genes were significant (Table 1; p<0.05). From these significant transcripts, 854 genes had a log fold change of greater than 1 while 837 genes had a log fold change of less than −1 (Figure 1B). Other RNA types were also significantly altered and these results are summarized in Table 1.

**Figure 1:**
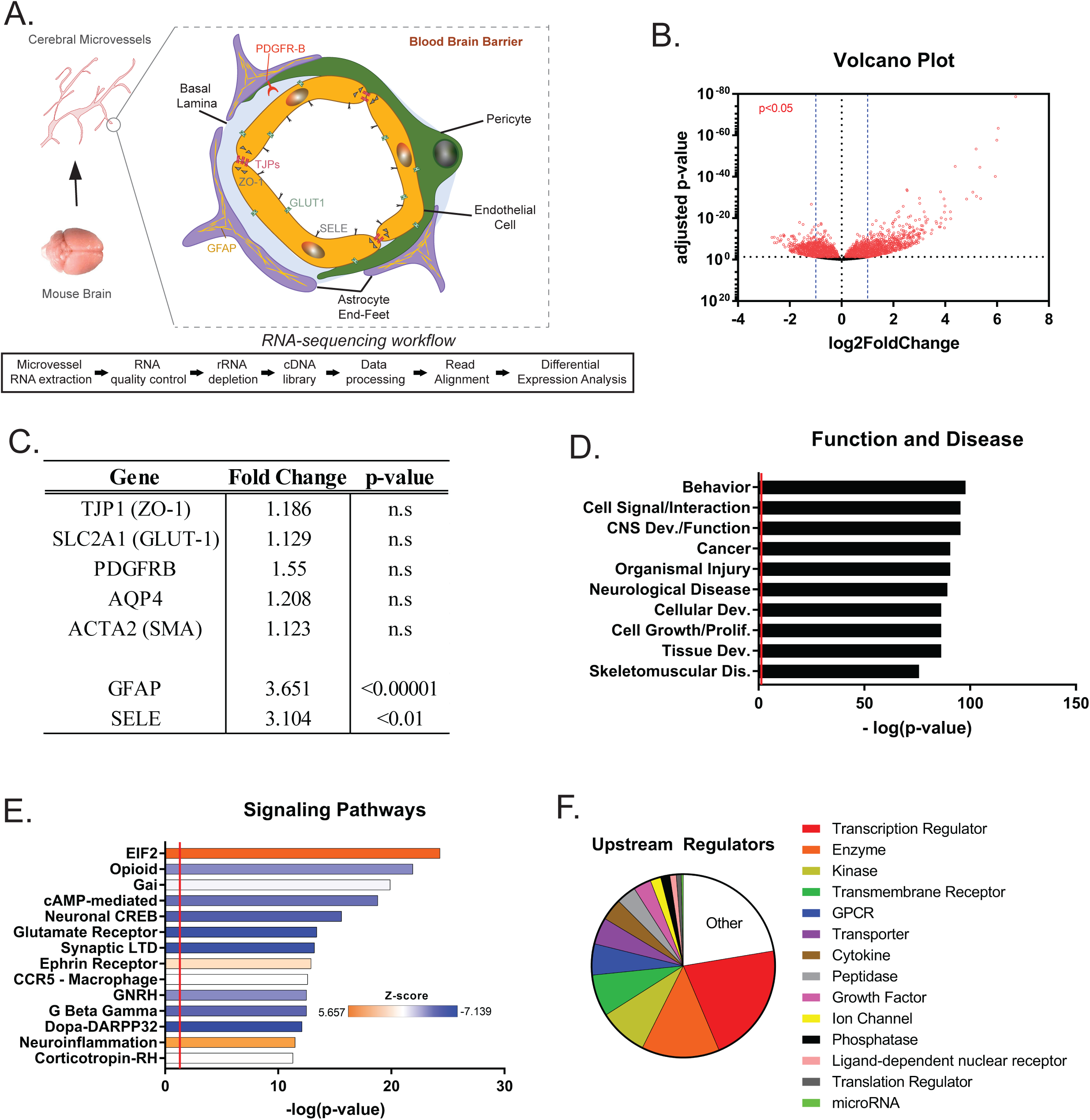
Transcriptomic changes in cerebral microvessels after tMCAO. A) Graphical summary of RNA-sequencing workflow and microvessel cell composition. B) Volcano plot of differential gene expression in microvessels after tMCAO. Red dots represent significantly altered genes (p<0.05). C) Table of microvessel component gene changes and neuroinflammation/endothelial activation gene changes. D) Bar plot of Disease and Function analysis generated by IPA. E) Bar plot of Canonical Signaling Pathway analysis generated by IPA. Prediction Z-score is overlaid on the bars with orange representing predicted pathway activation and blue representing predicted inhibition. F) Pie chart of predicted upstream regulators altered after tMCAO generated by IPA.

**Table 1:** Table of general overview of RNA-sequencing dataset from mouse microvessels after tMCAO.

| Transcriptome | Classification | Number |
| --- | --- | --- |
| mRNA | Total | 18491 (32129) |
|  | Significant Changes (p <0.05) | 6291 |
|  | > 1 Log2FC increase (p<0.05) | 854 |
|  | < -1 Log2FC decrease (p<0.05) | 837 |
| Noncoding RNA | Total | 13638 (32129) |
|  | Significant Changes (p <0.05) | 554 |
|  | lncRNA | 151 |
|  | Antisense RNA | 76 |
|  | miRNA | 14 |
|  | Others (Pseudogenes, SnoRNA, ScaRNA, etc) | 313 |

Cerebral microvessels are composed of endothelial cells, pericytes, and astrocytic end foot processes (Figure 1A). In order to examine the extent to which the cellular composition of the microvessels was altered after tMCAO, gene expression markers of these cell types were assessed (Figure 1C). In the RNA-sequencing dataset, the expression levels of these cell-identity markers remained insignificantly changed, whereas markers of neuroinflammation including glial fibrillary acidic protein (Gfap) and e-selectin (Sele) were significantly induced in microvessel preparations after tMCAO (Figure 1C). Similarly, qPCR validation of cell markers for pericytes (platelet derived growth factor receptor-beta (Pdgfr-beta) and CD45), astrocytes (aquaporin 4 (Aqp4)) and endothelial cell markers (zona occludens 1 (ZO-1)/tight junction protein 1 (Tjp1)) resulted in insignificant differences between sham and tMCAO microvessels (Supplemental Figure 2A-D).

**Figure 2:**
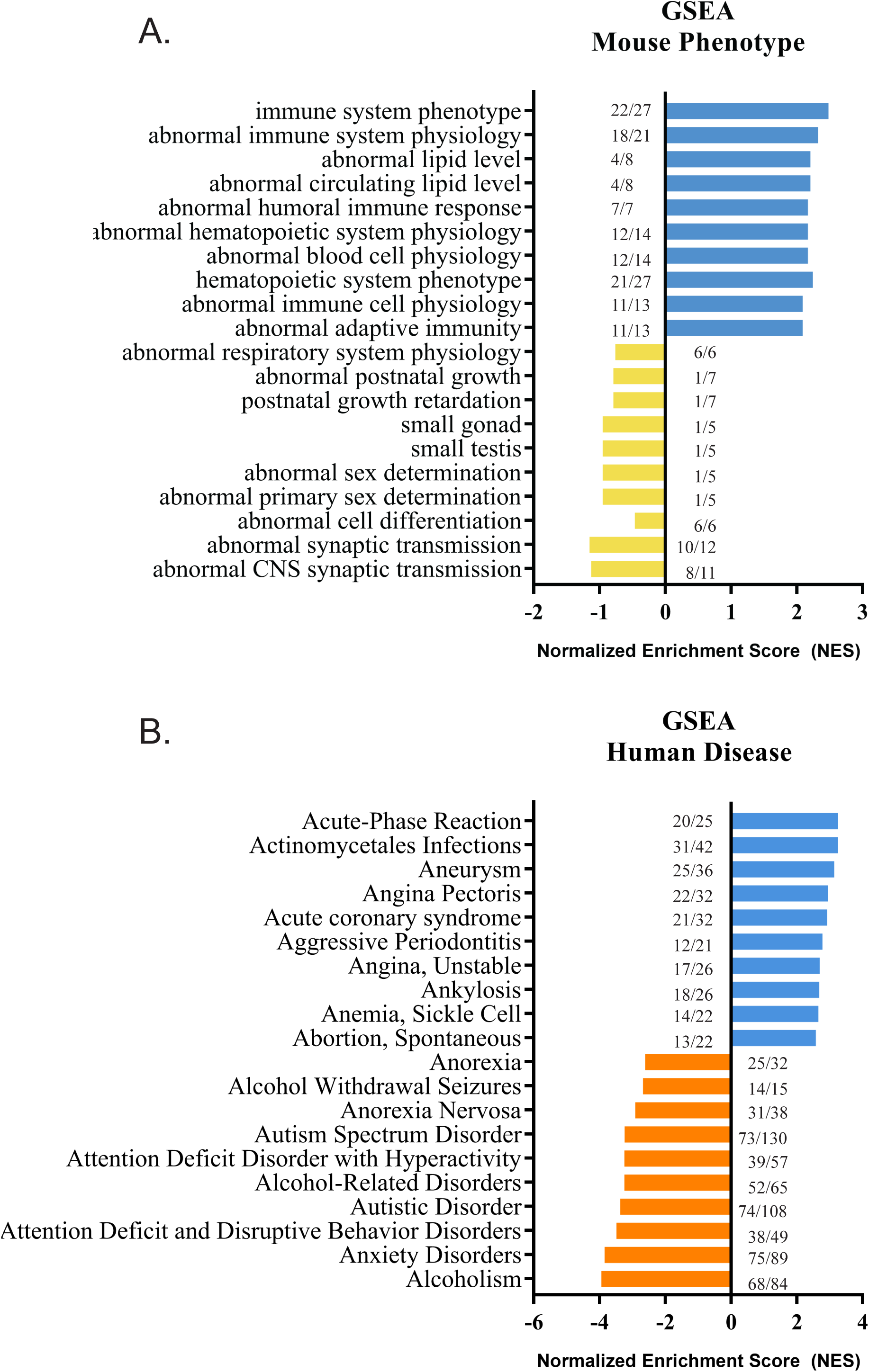
Translational relevance of microvessel-associated transcriptome changes. A) GSEA analysis of significant differentially-expressed genes in microvessels after tMCAO using a mouse phenotype database through WebGestalt. Ratios reflect number of genes in our dataset over the size of the *a priori* defined dataset. Blue bars represent a positive enrichment with an false discovery rate (FDR) of <0.05 and yellow bars represent a negative enrichment with an FDR of >0.05 but a p-value of <0.05. B) GSEA analysis of significant differentially expressed genes in microvessels after tMCAO using a human disease database (GLAD4U) through WebGestalt. Ratios reflect number of genes in our dataset over the size of the *a priori* defined dataset. Blue bars represent a positive enrichment with an FDR of <0.05 and orange bars represent a negative enrichment with an FDR of <0.05.

To identify the functional consequences of gene expression changes after tMCAO, we performed downstream effects (DE) analysis on a subset of the 6,291 significantly altered genes (with a log2 fold change (log2 FC) cut off of < - 0.5 and > 0.5) using Ingenuity Pathway Analysis (IPA). This DE analysis included Disease and Function, Canonical Pathway, and Upstream Regulator analyses that were used to identify predicted alterations in various functional and molecular categories based on gene expression alterations. Disease and Function analysis predicted alterations in various biological processes such as behavior, cell-to-cell signaling and interaction, central nervous system development and function, cancer, organismal injury, and neurological diseases (Figure 1D). Canonical Pathway analysis predicted positive pathway activation in eukaryotic translation initiation factor 2 (EIF2) and neuroinflammation signaling (Figure 1E). In contrast, predicted inhibited pathways shared downstream alterations related to G-coupled protein receptor (GPCR) signaling (ex: Gai, G Beta Gamma, and cAMP-mediated signaling) and calcium signaling (ex: CREB, Dopa-DARPP32, and Glutamate Receptor signaling) (Figure 1E). Due to the strength of predicted EIF2 activation indicating a potential halt in protein translation, we also examined the endoplasmic reticulum (ER) stress pathway. ER stress was predicted to be increased with implications for apoptosis (increased transcripts of caspase 3 and 7 (Casp3/7) and tumor necrosis factor (TNF) receptor associated factor 2 (Traf2)) along with decreased protein translation (Eif2). Further, we also examined the predicted upstream regulators affected after tMCAO (Figure 1E). The top five predicted upstream regulators were a part of following molecular categories: transcriptional regulators (21.25%), enzymes (13.75%), kinases (8.63%), transmembrane receptors (7.38%), and GPCRs (5.5%).

These results highlight the variety of molecular changes in the microvasculature after tMCAO. Neuroinflammation in these microvessel preparations is associated broad changes in ER functioning, calcium, and GPCR signaling that is predictive of both transcriptomic and metabolic changes at the cellular level after tMCAO.

### Translational relevance of microvessel-associated transcriptome changes

As many transcriptomic alterations were observed in microvessel preparations after tMCAO, we next sought to specifically examine how these changes related to human disease. To this end, the WebGestalt tool was used to perform gene set enrichment analysis (GSEA) of *a priori* defined gene sets related to mouse phenotype and human disease. For mouse phenotypic analysis, GSEA was performed using the Mammalian Phenotype Ontology database from Jackson labs (MP:0000001; Figure 2A). From this analysis, we observed that genes involved in immune response and abnormal lipid levels (ex: TNF super family member 9 (Tnfsf9), C-C motif chemokine ligand 2 (Ccl2), and complement component 2 (C3)) were positively enriched while genes involved in postnatal development and central nervous system transmission were negatively enriched (Supplemental File 1). To further understand the translational relevance of this data, we performed an additional GSEA to identify enriched classes of genes involved in human disease using the GLAD4U database (Figure 2B). As expected, these same genes (ex: Ccl2 and other TNF-family members) were positively enriched with human diseases with a large inflammatory or hypoxic component such as Acute-Phase Response, infections, and aneurysm (Supplemental File 1). Of interest, markers of endothelial cell activation including matrix metalloproteinase 9 (Mmp9), Intercellular Adhesion Molecule 1, (Icam1), and TNF family members were also positively enriched in these disease categories. Interestingly, genes involved with more neurological diseases and function were significantly and negatively enriched.

Additional analyses were completed using the Disgenet and OMIM databases for human disease ontology and produced comparable results (Supplemental Figure 3). These results highlight the enrichment of inflammatory and neurological disease categories in microvessel-associated gene expression changes after tMCAO. Additionally, these comparisons provide further support for the relevance of our microvessel model in capturing these aspects of stroke pathology.

**Figure 3:**
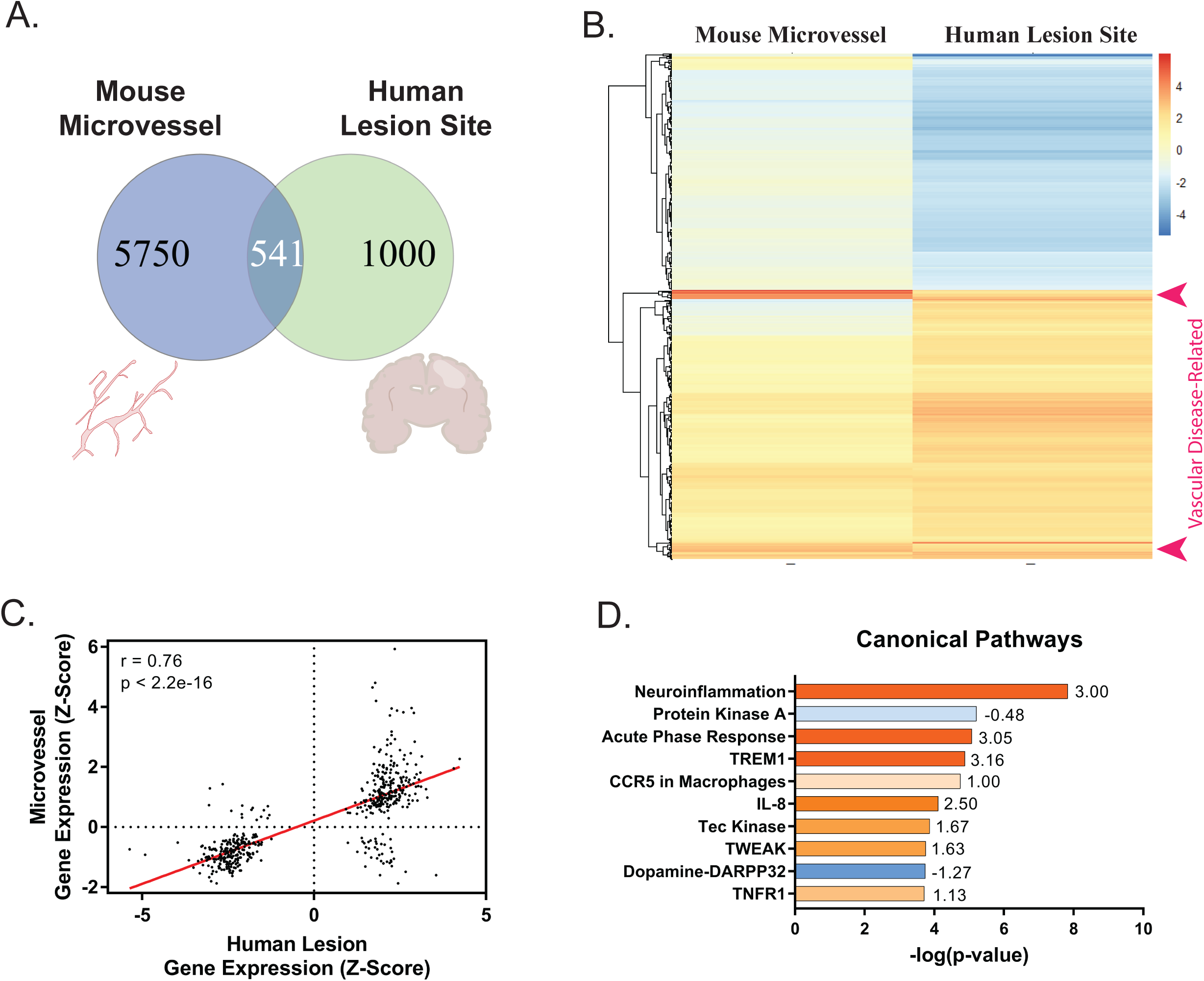
Comparison of mouse microvessels to human lesion samples after ischemia. A) Venn diagram of gene expression comparisons between mouse microvessels and human lesion-sites. B) Heatmap of the differential gene expression Z-scores of 541 shared genes ranked by k-means clustering (seed = 25). C) Pearson correlation of transcript expression Z-scores between mouse microvessels and human lesion-sites (r = 0.76; p < 2.2e-16). D) Bar plot of Canonical Signaling Pathway analysis of 541 shared genes generated in IPA. Predicted activation and inhibition is overlaid on the bars with the Z-score printed adjacent.

### Comparison of mouse microvessels to human lesion samples after ischemia

In an effort to understand the specific changes in gene expression relevant to human stroke, we conducted a direct comparison of our microvessel dataset with a publicly-available dataset containing RNA-sequencing data of lesion-site samples from patients post-stroke. Further information on this dataset is summarized in Supplementary Table 1. Human samples in this dataset included tissue from cortical lesion sites of patients with advanced age (67-74 years) and a history of non-fatal ischemia within the last 5 years (GSE56267).

Initial comparison of these datasets reveal 541 shared genes, accounting for 8.6% of the microvessel dataset and 35.1% of the human dataset (Figure 3A). A heatmap of the Z-scores of the log2FC for these genes is represented in Figure 3B. To further investigate the how these 541 genes may have related one another, we performed k-means clustering to divide these genes into 5 distinct clusters (Supplemental Figure 4A and 4B). As could be expected, these distinct clusters were associated with distinct aspects of the shared gene expression patterns between the mouse microvessels and human lesion sites. For example, while clusters 3 and 4 both contained genes involved in inflammation, cluster 3 contained general markers of this response (including genes transcribing TNF-family members, NF-kb, and chemokines) and cluster 4 contained 18 vascular disease specific genes involved in inflammation (including heme oxygenase 1 (Hmox1), serpin e family member 1 (Serpine1), pentraxin-related protein 3 (Ptx3), tissue inhibitor of metalloproteinase 1 (Timp1), and CD44) (Figure 3B, Supplemental Figure 4C). Cluster 5 contained negatively enriched genes implicated in neurotransmission and metabolism. Additionally, these clusters corresponded to previously defined diseases with separate clusters containing genes enriched for inflammation (cluster 3), central nervous system (cluster 5), and vascular disease (cluster 4) (Supplemental Figure 4D). These results highlight the functional compartmentalization of the gene alterations shared between our mouse microvessel model and human lesion sites.

**Figure 4:**
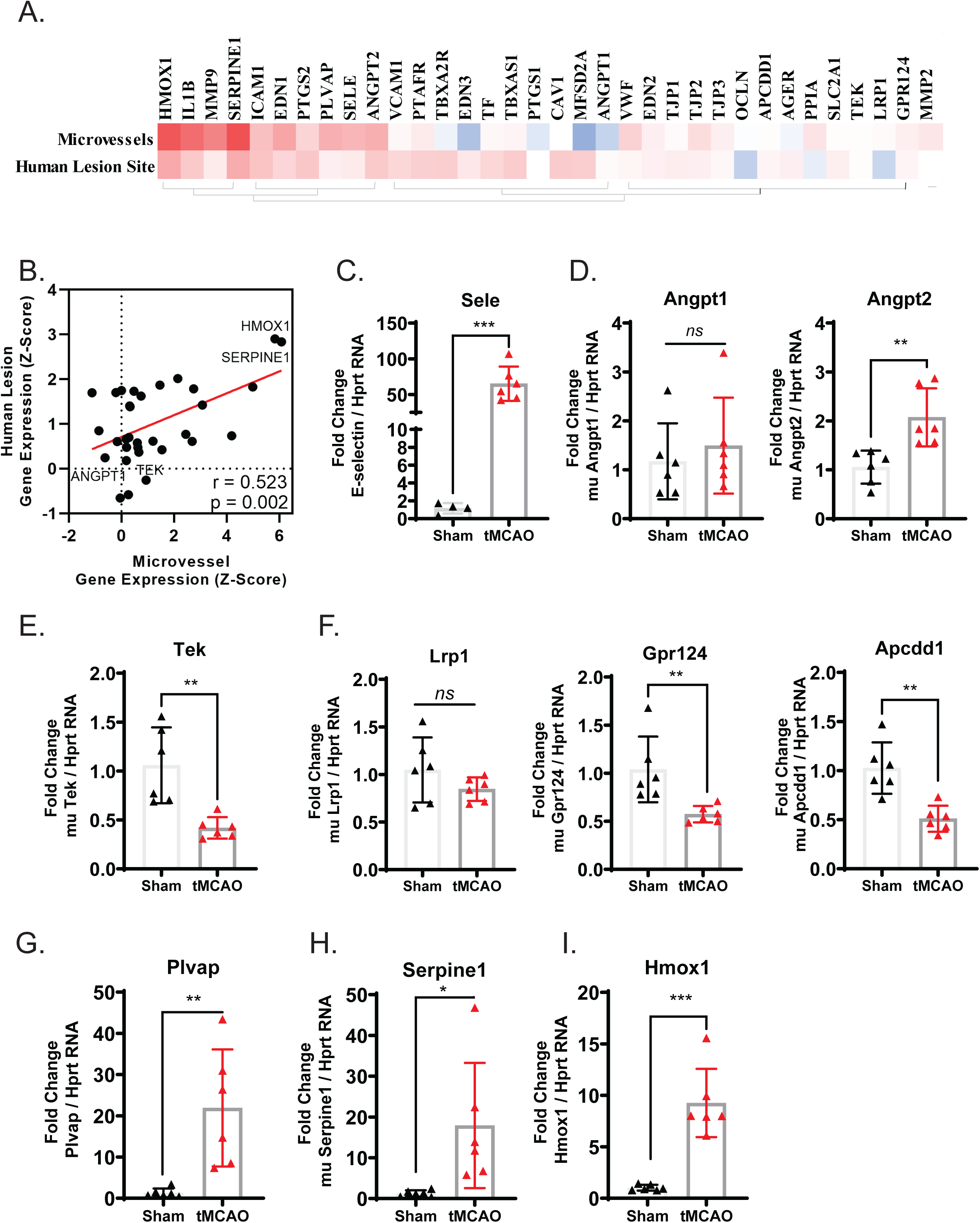
Ischemia-induced endothelial cell activation alters BBB integrity in cortical microvessels. A) Heatmap of genes relevant to endothelial cell activation and BBB integrity and function ordered by hierarchical clustering. B) Pearson correlation of transcripts represented in 4A between RNA-sequencing data of mouse microvessels after tMCAO and human stroke lesion-sites (r = 0.52; p < 0.01). QPCR validation of transcripts involved in BBB function including: C) endothelial cell activation, D-E) angiopoietin ligands and receptor, F) WNT signaling, G-I) vascular permeability and endothelial function. (*p<0.05; **p<0.01; ***p<0.001; ns = not significant).

Further, the 541 genes shared between the human lesions and microvessels were expressed in a highly similar manner (Figure 3C; r = 0.76). Functional analysis with of these genes predicted multiple canonical pathways involved in neuroinflammation to be significantly upregulated while g-coupled protein receptor (Protein Kinase A) and calcium dependent signaling processes (Dopa-DARPP) were predicted as significantly suppressed (Figure 3D). As expected, we then see that genes involved in Neuroinflammation (r = 0.86, p < 0.0001, n = 27) and Protein Kinase A (r = 0.87, p < 0.00011, n = 25) were similarly expressed across both datasets (Supplemental File 2). While gene expression in the Dopa-DARPP category as a whole was not correlated (r = 0.30, p = 0.31, n = 13), if we removed the dopamine specific genes to highlight the calcium and g-protein coupled receptor signaling components we saw a dramatic correlation of expression between mouse microvessels and human lesion sites (r = 0.98, p = 0.0006, n = 6). These removed genes included voltage-gated potassium channels and DARPP32-specific targets not relevant to microvessel cell types (Supplemental File 2).

These findings further highlight the conserved alteration of these inflammation and metabolic pathways in ischemic human lesions and mouse microvessels. Given these results, we interpret that the relevance of these microvessel-associated changes to human pathophysiology may lie largely in the disruption of the BBB.

### Ischemia-induced endothelial cell activation alters transcripts related BBB function in cortical microvessels

In order to more closely examine BBB-related transcriptomic alterations after tMCAO and stroke, we included 34 genes related to endothelial cell activation and BBB function and maintenance (Figure 4A). Expression of these genes was significantly correlated between mouse microvessels and human lesion-sites after stroke (Figure 4B). Molecules involved in maintaining BBB and endothelial cell function are exciting potential targets in neurovascular disease and we observed that several of these genes (including Hmox1, Serpine1, CD44, gardner-rasheed feline sarcoma viral (Fgr), and vitamin D receptor (VDR)) were similarly induced after tMCAO in mice and chronically post-stroke in humans (Figure 4B). Further, endothelial cell activation indicated by robust Sele induction was validated in qPCR (Figure 4C).

Subsequent validation of some of these translationally-relevant targets revealed novel genes altered in microvessel preparations after tMCAO. Of interest, transcriptional changes in the family of angiopoietins 1/2 (Angpt1/2) and their receptor, tyrosine-protein kinase receptor (TEK), were validated by qPCR. While the transcriptional difference in Angpt1 was insignificant, Angpt2 transcript was significantly induced in the microvessel preparations post-tMCAO (Figure 4D). Tek, the gene that transcribes the Tie2 receptor for Angpt1/2, was also down regulated after tMCAO (Figure 4E).

In addition, we also analyzed some of critical players of wingless (WNT) signaling in the context of BBB maintenance and function. qPCR-based validation revealed significant down regulation in the mRNA levels of g-protein coupled receptor 124 (Gpr124) and adenomatosis polyposis coli down-regulated 1 (Apcdd1) with an insignificant effect on low density lipoprotein receptor-related protein 1 (Lrp1) mRNA levels in tMCAO microvessels (Figure 4F).

Additional analysis of the 541 shared genes between mouse microvessels and human lesion-sites revealed 68 genes that were existing drug targets (Supplemental Figure 5). This list contained several endothelium-associated genes, including Hmox1 and Serpine1, which were robustly shared between the two datasets. The strong induction of these two specific transcripts was validated in microvessels after tMCAO by qPCR (Figure 4G-I). Hmox1 and Serpine1 are of high interest as possible novel and druggable targets for the microvasculature and have had promising experimental and clinical outcomes for stroke (Ref). Thereafter, we sought to validate the enrichment of the selected BBB maintenance and function-associated targets in the microvessel preparations. We performed qPCR quantification of Angpt1/2, Tek, Lrp1, Gpr124, Apcdd1, plasmalemma vesicle-associated protein (Plvap), Serpine 1 and Hmox1 transcripts in the total RNA extracted from whole brain, cortex, and isolated microvessels of naïve mice. These genes were significantly enriched in the microvessels when compared to whole brain and cortex samples suggesting their potential relevance in microvessel-specific functions associated with the maintenance and proper functioning of the BBB (Figure 5A-E).

**Figure 5:**
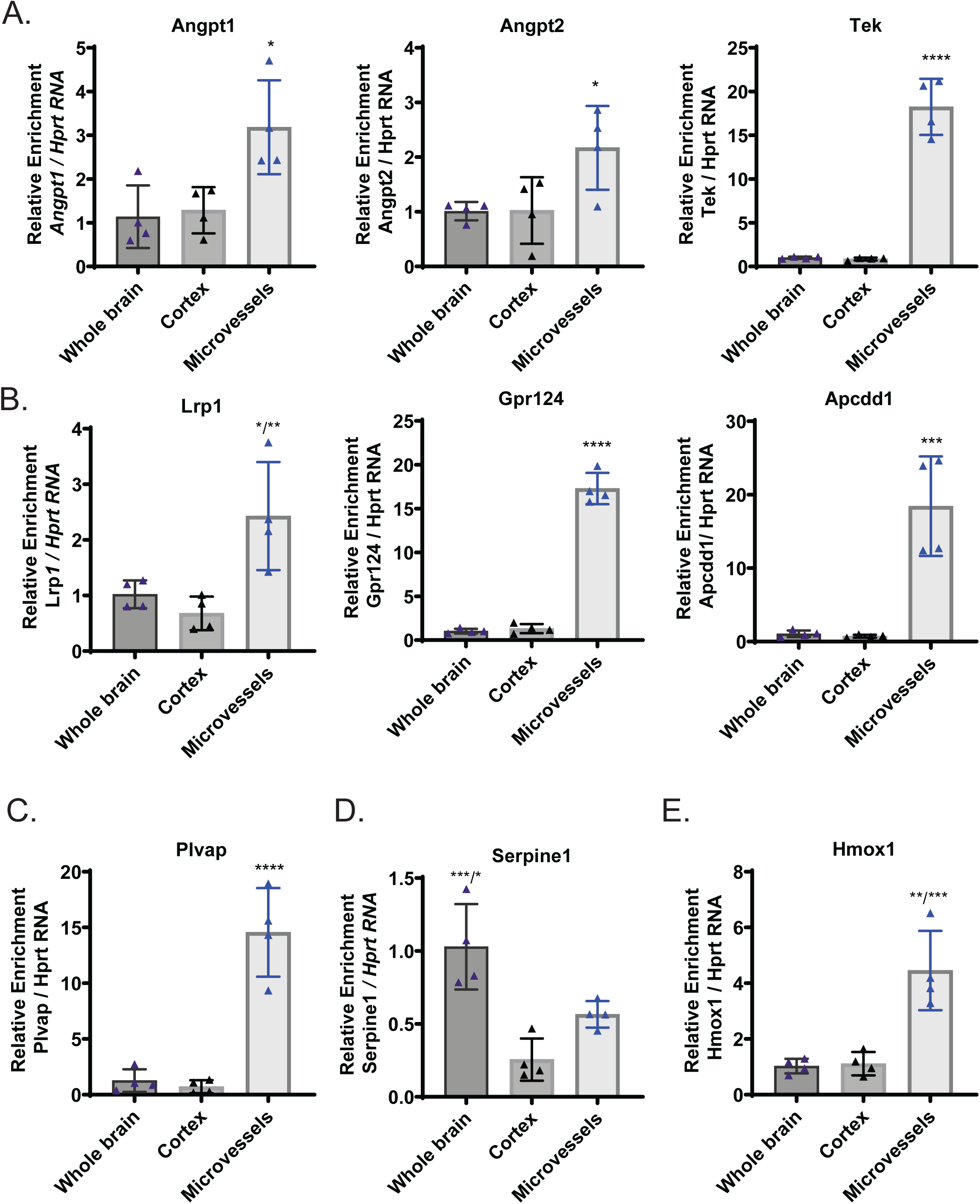
Target genes related to BBB function are enriched in cortical microvessels. A-E) qPCR validation of the enrichment of selected BBB target genes in isolated mouse microvessels compared to whole brain and cortex. (*p<0.05; **p<0.01; ***p<0.001); ****p<0.0001; ns = not significant).

These results support and emphasize the importance and relevance of the neurovasculature to stroke and highlight the potential of targeting microvessel and endothelial cell function in humans. While many of these transcript alterations have been described to play a role in BBB maintenance and function, how these genes are regulated upon injury was still previously unknown. The observed conservation of pathways related to endothelial cell activation and BBB integrity between tMCAO microvessels and human brain samples after stroke strongly support the principle that these changes at the neurovasculature level are crucial in sustained pathogenesis after ischemia.

### Ischemia-associated changes in S1P metabolism and receptor signaling in cerebral microvessels

As a novel contributor of endothelial cell activation and BBB function, we decided to examine transcriptomic changes in mediators of sphingolipid metabolism. A succinct summary of this pathway with our qPCR results is represented in Figure 6A and a full heatmap containing 125 genes involved in sphingolipid metabolism (GO: 0006665) is available in Supplemental Figure 6A. These 125 genes were expressed in a similar manner between mouse microvessels and human lesion sites (Figure 6B).

**Figure 6:**
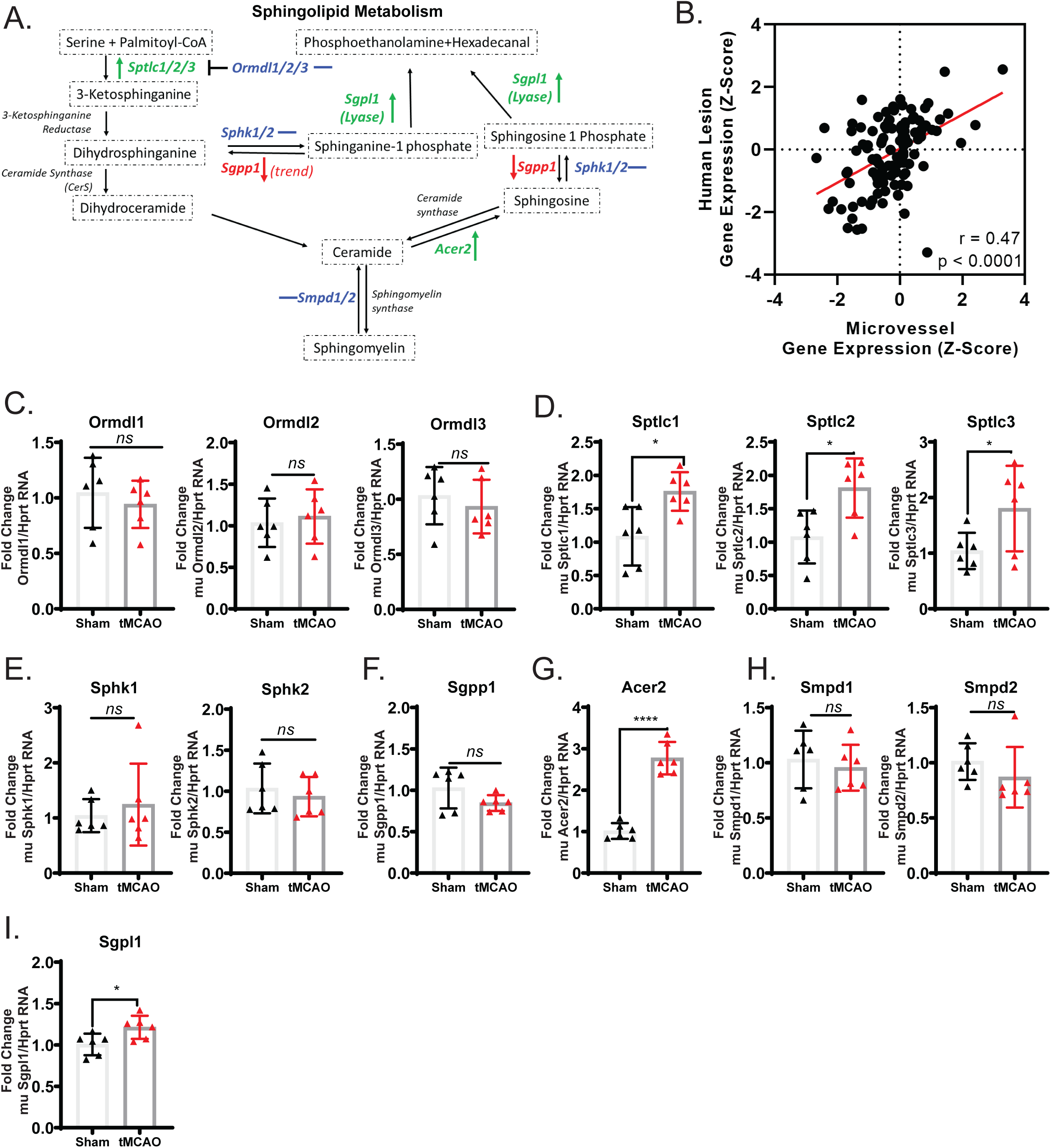
Ischemia-associated changes in S1P metabolism genes in cerebral microvessels. A) Simplified summary of sphingolipid metabolism with qPCR results included (green: upregulated; blue: no change; red: downregulated). B) Pearson correlation of transcript expression from mouse microvessels and human lesion-sites including genes from sphingolipid metabolic processes (GO: 0006665). qPCR validation of transcripts involved in sphingosine metabolism separated into functional groups of genes for C) sphingolipid biosynthesis regulators (Ormdl1/2/3), D) serine palmitoyltransferases (Sptlc1/2/3), E) sphingosine kinases (Sphk1/2), F) S1P phosphatase (Sgpp1), G) alkaline ceramidase 2 (Acer2), H) sphingomyelinases (Smpd1/2), and I) S1P lyase (Sgpl1). (*p<0.05; ****p<0.0001; ns = not significant).

Specifically related to the synthesis of sphingosine, we conducted qPCR validation of selected targets involved. While there were no significant changes in the serine palmitoyltransferase (SPT) inhibitors, ORMDL sphingolipid biosynthesis regulator 1/2/3 (Ormdl1/2/3), after tMCAO there was a modest and significant increase in the SPT genes themselves, serine palmitoyltransferase long chain base subunit 1/2/3 (Sptlc1/2/3) (Figure 6C-D). Further, there were no observed differences in the transcripts for sphingosine kinases 1/2 (Sphk1/2) or S1P phosphatase 1 (Sgpp1) after tMCAO (Figure 6E-F). Interestingly, there was a significant elevation of alkaline ceramidase 2 transcript (Acer2) suggesting the potential for an increase in conversion of ceramide to sphingosine (Figure 6G). Additionally, while there was a modest increase in S1P lyase 1 (Sgpl1) transcript and no remarkable differences in sphingomyelin phosphodiesterase (Smpd1/2) transcript expression (Figure 6H-I). To better understand the tissue specificity of S1P metabolism-associated genes we conducted a qPCR-based relative enrichment analysis between the total RNA extracted from whole brain, cortex tissue, and cortical microvessels of naïve mice. The genes Orndl1/3, Sptlc1/2/3, Sphk1/2, Sgpp1, Smpd1/2 and Sgpl1 were significantly enriched in the microvessel preparations when compared to whole brain and cortex samples (Figure 7A-G). Importantly, levels of Acer 2 were significantly and robustly enriched in the microvessel fraction compared to cortex and whole brain (Figure 7E). In addition, the mRNA levels of Ormdl2 seem modestly but insignificantly abundant in the microvessel preparations when compared to whole brain tissue (Figure 7B).

**Figure 7:**
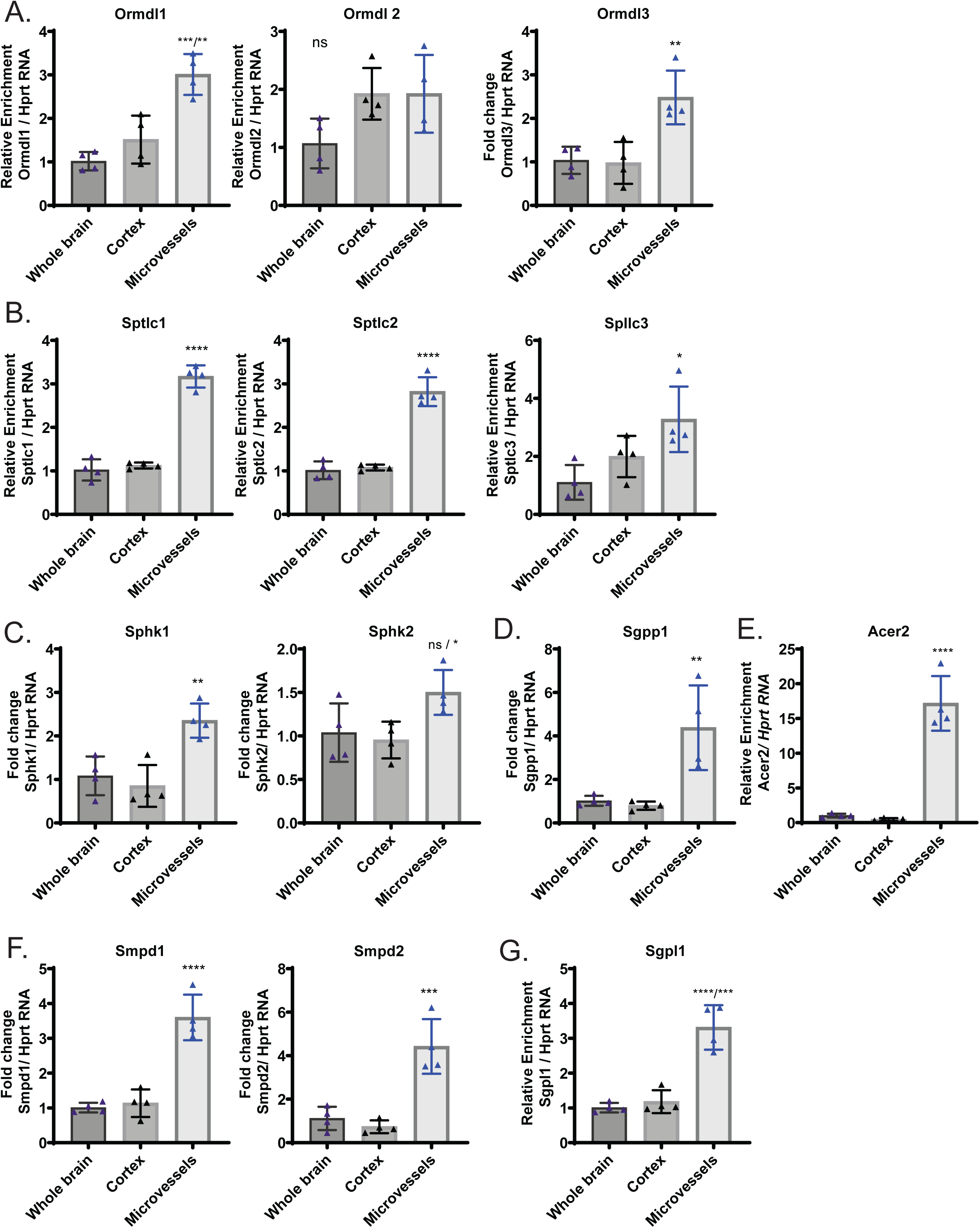
Specific target genes related to S1P metabolism are enriched in cortical microvessels. A-G) qPCR validation of the enrichment of sphingolipid metabolism target genes in isolated mouse microvessels compared to whole brain and cortex. (*p<0.05; **p<0.01; ***p<0.001; ****p<0.0001; ns = not significant).

Additional qPCR validation of mRNA transcribing S1P’s cognate receptors that mediate sphingolipid signaling was conducted. Significant changes in S1pr1/2/4 were observed in microvessels after tMCAO. Of note, S1pr2 was the most robust and significant receptor alteration, in line with previous data from our lab (Supplemental Figure 6D). Supplmental analysis of RNA-sequencing data with differential gene expression data overlaid on the sphingosine receptor signaling pathway from IPA is available in Supplemental Figure 6C. This data is further reflected in a heat map to compare these changes in mouse microvessels and human lesion-site samples (Supplemental Figure 6B).

Taken together, we highlight novel transcriptomic alterations in sphingosine metabolism that provide additional support to the investigation of this pathway in stroke. This data suggests the importance of the sphingolipid metabolic axis in the cerebral microvessels that might be essential in the regulation of BBB function. Specifically, Acer2 has emerged as a potential target enriched in the microvessels that warrants further investigation. As a major player in ceramide metabolism, this induction is associated with neuroinflammation-associated endothelial cell activation.

## MATERIALS AND METHODS

### Animals and ischemic model

All animal experiments were approved by Weill Cornell Institutional Animal Care and Use Committee. C57/BL6/J (8-10 weeks, male) mice are subjects to transient middle cerebral artery occlusion (tMCAO). During tMCAO surgery, mice were deeply anesthetized with Isoflurane and body temperature was maintained at 37 °C with self-regulating heating pad. The flow of middle cerebral artery was temporally occluded with silicon coated intraluminal suture (Doccol, 602245PK10Re) for 60 minute and reperfused by withdrawing a suture, as previously described [26]. Animals which unable to walk straight 24 hours after surgery were used for microvessel isolation.

### Microvessel isolation

The cerebral microvessels were isolated as previously described[25]. To minimize cell activation, all procedures were conducted in a cold room. Ipsilateral cortices were homogenized with MCDB131 medium (Thermo Fisher Scientific, 10372019) with 0.5% fatty acid free BSA (Millipore Sigma, 126609). The homogenate was centrifuged at 2000 g for 5 minutes at 4°C. The pellet was suspended in 15% dextran (molecular weight ∼70 kDa, Millipore Sigma, 31390) in PBS and centrifuged at 10000 g for 15 minutes at 4°C. The pellet was resuspended in MCDB131 with 0.5% fatty acid free BSA and centrifuged at 2000 g for 10 min at 4°C. The pellet contained the microvessels.

### RNA extraction from cerebral microvessels

Isolated cerebral microvessels were dissolved in appropriate amount of RLT lysis buffer to lyse the multicellular structures. Lysed samples were then loaded onto a column-based shredder (QIAshredder, Qiagen, Germany) and centrifuged (8000g for 2min at room temperature) to eluate the homogenized lysate. Thereafter, the lysate was processed using total RNeasy mini kit (Qiagen, Germany). In brief, the lysate was resuspended in 1:1 volume 70% ethanol and loaded onto extraction column and centrifuged (8000g for 2min) at room temperature. DNase-I digestion was performed on-column for samples with 1U of TURBO-DNase (Invitrogen, ThermoFisher Scientific) and incubated for 15min at room temperature. Then columns were washed with appropriate amount of wash buffer and finally total RNA was eluted using 25-20 µl of nuclease free water.

### Quality Check of total RNA extracted from cerebral microvessels

To ensure the quality of total RNA preparations from the cerebral microvessels, 200ng of total RNA were electrophoresed in 1% Agarose Gel at 80V for 30min at room temperature. Further, 100ng of total RNA preps for each sample were analyzed for A260/280 ratios using UV-visible spectroscopy in a microplate format (Varioskan, ThermoFisher Scientific) and subsequently confirmed with a final analysis using Bioanalyzer quantification (Genomics core facility, Weill Cornell Medicine).

### RNA-sequencing workflow

RNA integrity and other quality control measures were performed, as stated above, along with ribosomal RNA depletion before cDNA library preparation. cDNA libraries were made using the Illumina TruSeq Stranded mRNA Library Prep kit and were sequenced in a single lane with pair-end 101 bps on Illumina HiSeq4000 instrument. Cutadapt was used to trim low quality bases and adapters [27]. STAR was used to align raw sequencing reads to the mouse GRCm38 - mm10 reference genome[28] Raw read counts were calculated using HTseq-count[29]. Differential expression analysis was performed using DEseq2 package [30]. Volcano and bar plots along with pie chart were generated in GraphPad Prism.

### Gene Set Enrichment and Functional Prediction Analysis

The downstream analysis (including Disease and Function, Canonical Pathways, and Upstream Regulator analyses) reported in this study were generated through the use of IPA (QIAGEN Inc., https://www.qiagenbioinformatics.com/products/ingenuity-pathway-analysis)[31] Significant genes for downstream analysis in IPA were included with a cut-off of −0.5 and 0.5 log2 fold change (p<0.05). GSEA analysis was completed using the WebgestaltR package in R Studio with all significantly altered[32]. Mouse phenotype GSEA was completed using the Mammalian Phenotype Ontology database from Jackson Labs (MP:0000001; Accessed on 11/14/2018 by WebGestalt tool). Human Disease GSEA was completed using DisgeNET (Version 5.0, 05/28/2017), Online Mendelian Inheritance in Man (OMIM; https://www.omim.org/), and GLAD4U (Disease terms were downloaded from PharmGKB (Accessed 11/2018 by WebGestalt tool)) databases. Genes associated with individual disease term were inferred using GLAD4U.

### Dataset comparison

Data processing was completed using the R Studio statistical computing environment (http://cran.us.rproject.org). The Vennerable and limma packages in R Studio were used to compare datasets and generate a venn diagram [33]. The pHeatmap package was used to generate the gene expression heatmaps. Linear regression and Pearson’s correlation tests were used to measure associated gene expression between mouse microvessels and human lesion sites. Gene expression heatmaps with named genes were clustered in R with the hclust and kmeans packages and further generated in excel. All clustering was conducted with a seed set at 25.

### cDNA synthesis and quantitative real time PCR (qPCR)

100-200 ng of total RNA was used as template to prepare cDNA for each sample in the presence of Verso-Reverse Transcriptase and Random Hexamer Primers (Thermo Scientific, USA) at 42 °C for 30 min in a final reaction volume of 20µl. Then, the cDNA prep was diluted 5-fold with nuclease free water to obtain a working stock at a final volume of 100 µl. SYBR Green chemistry with Rox dye signal normalization (Quanta bio, MA, USA)-based quantitative PCR was performed to determine the relative expression levels of target gene. qPCR reactions were run either in duplicates or, triplicates on a 96 well plate with 2µl of cDNA as template in a final reaction volume of 10µl per well for each sample. Finally, the plate was loaded on to the ABI-7500 Sequence Detection System PCR machine (Applied Biosystems, USA). Amplification of target genes were performed using the following cycling conditions with initial denaturation at 95°C for 10min followed by 30cycles of 95 °C for 10s; 56 °C for 30s (annealing), extension at 72 °C for 30s with melt curve analysis at 60 °C for 30s at a linear ramp rate of 0.5 °C/s followed by acquisition at 0.5 °C intervals. First, the expression levels (Ct values) of target gene were normalized to expression levels of Hprt rRNA as ΔCt values. For Sham and Stroke samples, the relative expression levels of target gene were then expressed as ΔΔ Ct values by comparing the ΔCt values of each sham to the average ΔCt of the sham group. Fold change in target gene expression was calculated by comparing the 2-Δ Ct values of the stroke samples with that of sham samples. A detailed list of primers sequences for the target genes used in this study is provided in table 2.

**Table 2:**
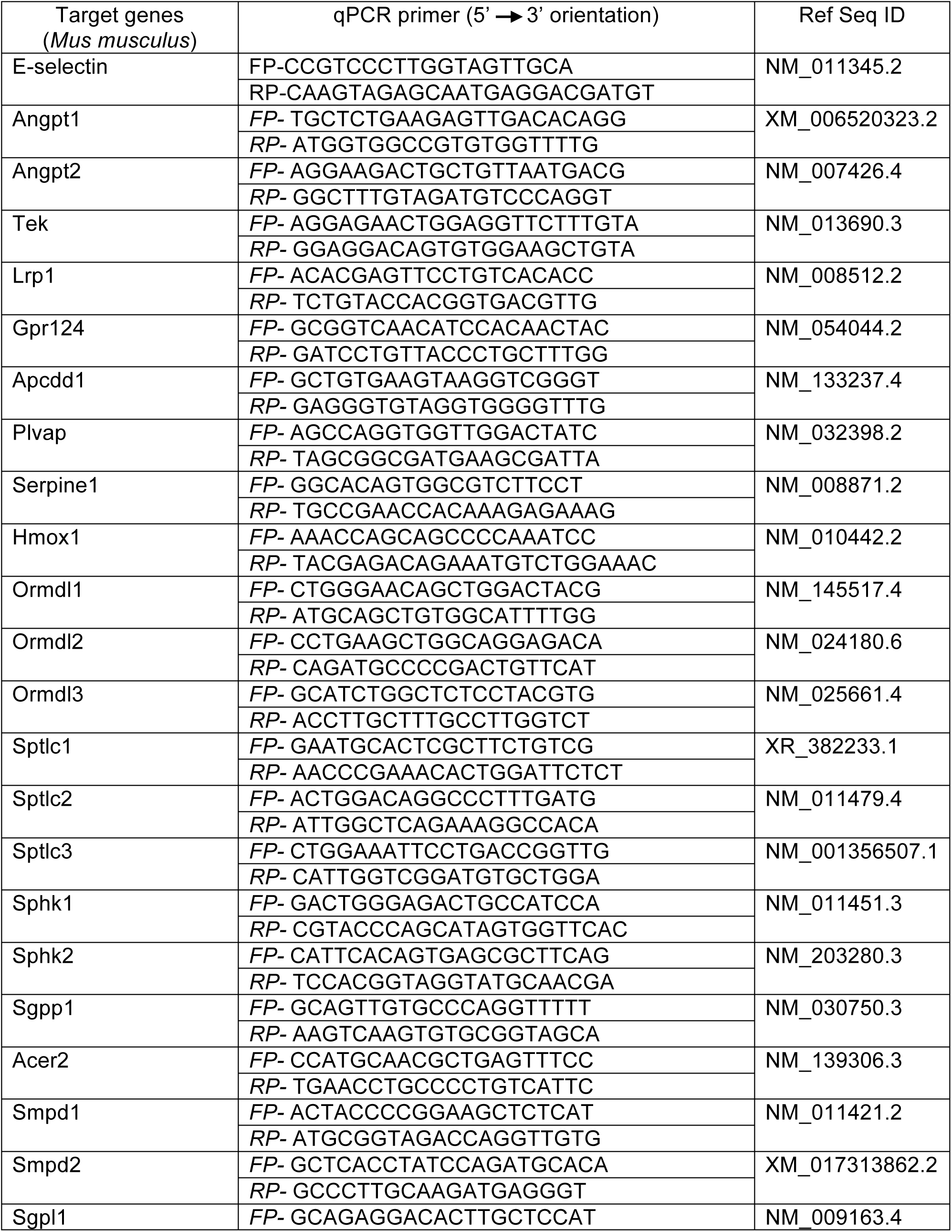
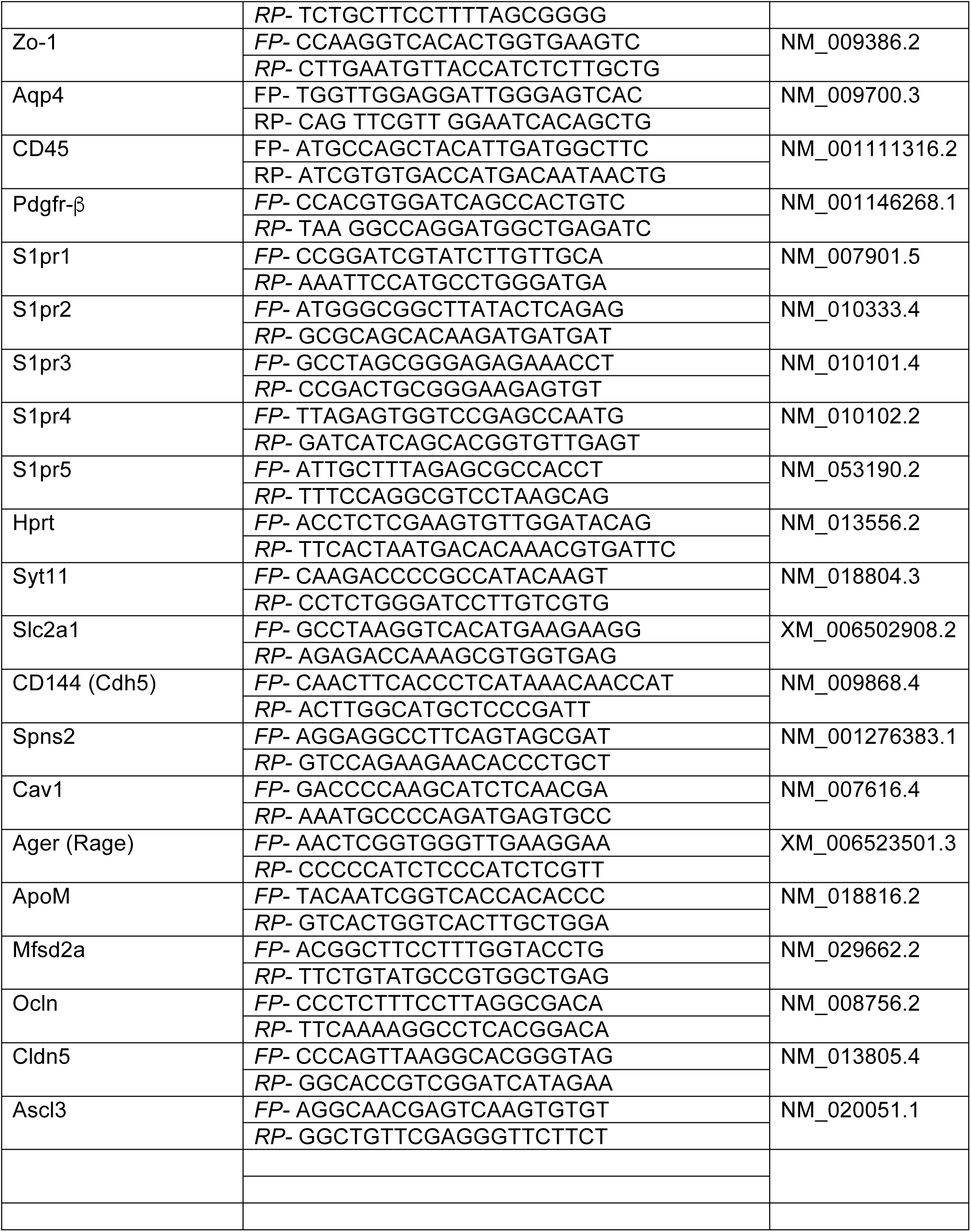
List of primer sequences used in qPCR experiments.

### Data Availability

The human lesion site data used in these analysis is previously published and available in the NIH GEO Omnibus under accession number GSE56267[34].

## ACKNOWLEDGEMENTS

This work was supported by internal funds provided by the Department of Pathology and Laboratory Medicine, Weill Cornell Medicine and American Heart Association Grant-in-Aid 12GRNT12050110, NIH HL094465 and Leducq Foundation grants to TS. AI was partially supported by LeRoche foundation and Tri I TDI.

The authors declare no competing financial interests.

---

**Supplemental Figure 1:**
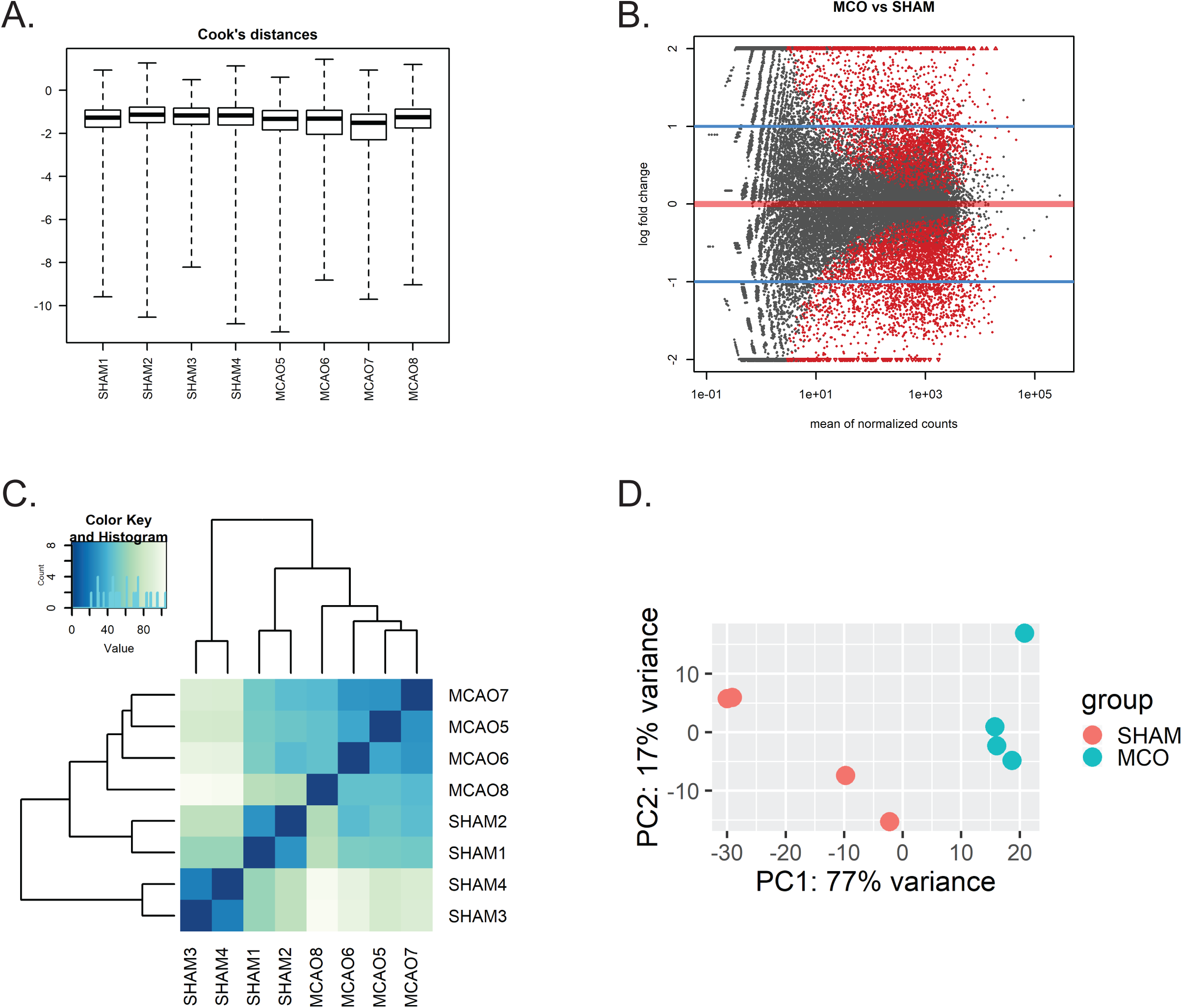
A) Cook’s distances of all samples used in differential expression analysis. B) MA plot depicting global view of the relationship between the expression change between sham and tMCAO samples (log2 FC, M) and the average expression strength of the genes (mean of normalized counts, A) with significance threshold overlaid (red = p < 0.05). C) Heatmap of Euclidean sample-to-sample distance matrix across all samples. D) Principal Component Analysis (PCA) of sham and tMCAO (MCO) samples.

**Supplemental Figure 2:**
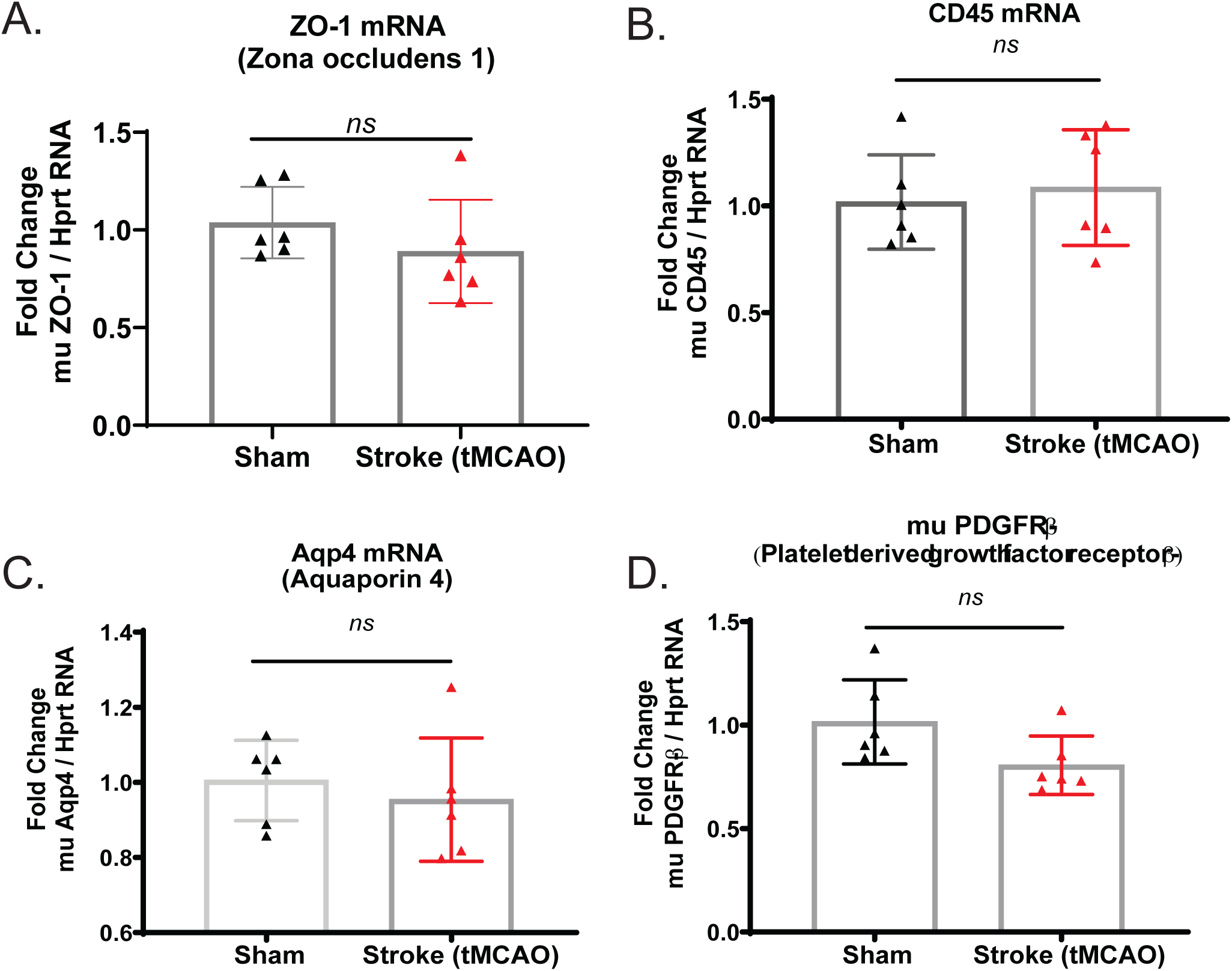
qPCR validation of cell component transcripts in microvessels isolated from sham and tMCAO mice. (**p<0.01; ***p<0.001); ns = not significant).

**Supplemental Figure 3:**
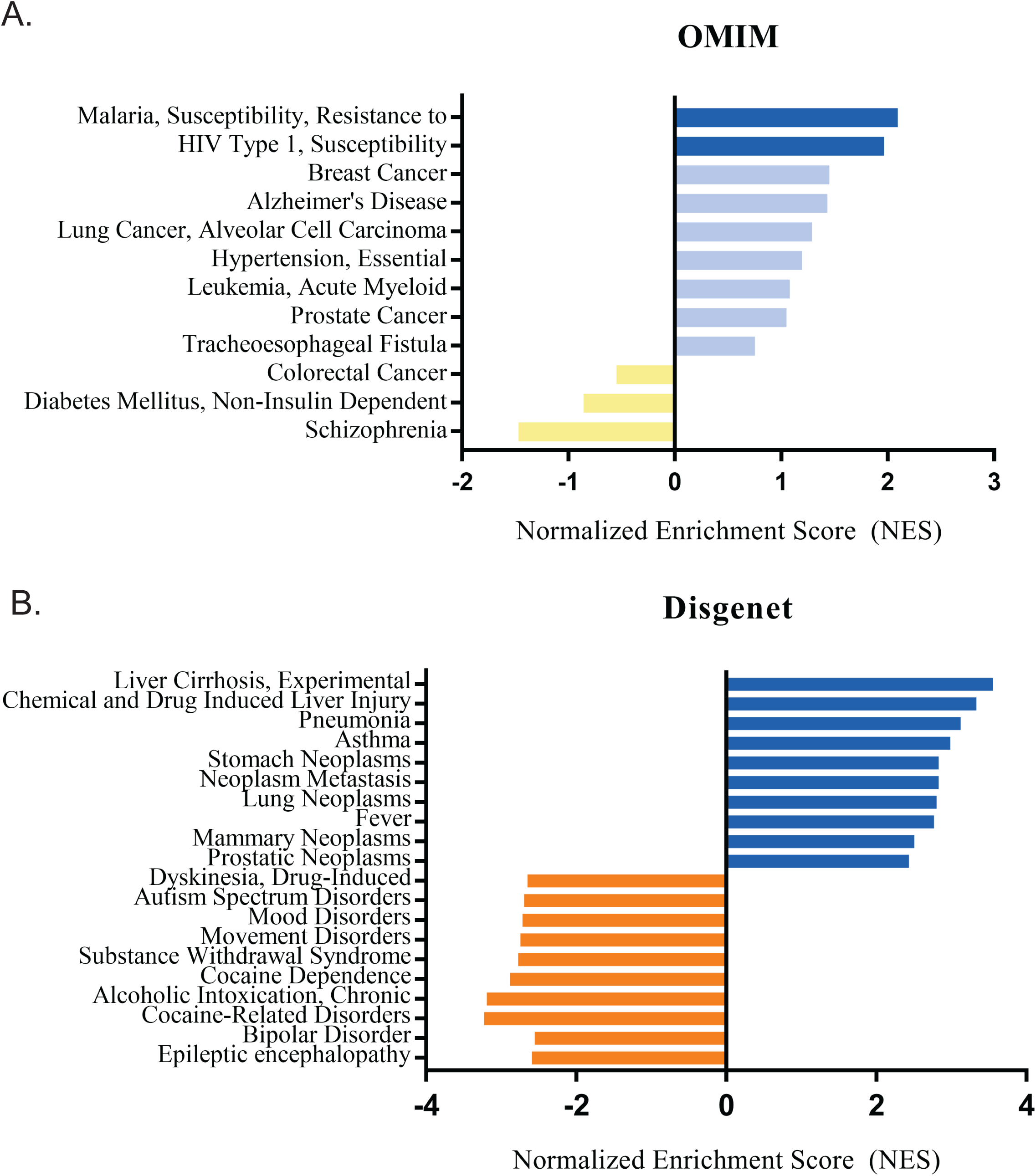
A) GSEA analysis of significant differentially expressed genes in microvessels after tMCAO using a human disease database (OMIM) through WebGestalt. B) GSEA analysis of significant differentially expressed genes in microvessels after tMCAO using a human disease database (Disgenet) through WebGestalt. Dark blue bars represent a positive enrichment with an FDR of <0.05, light blue bars represent a positive enrichment with an FDR of >0.05 but a p-value of <0.05, yellow bars represent a negative enrichment with an FDR of >0.05 but a p-value of <0.05, and orange bars represent a negative enrichment with an FDR of <0.05.

**Supplemental Figure 4:**
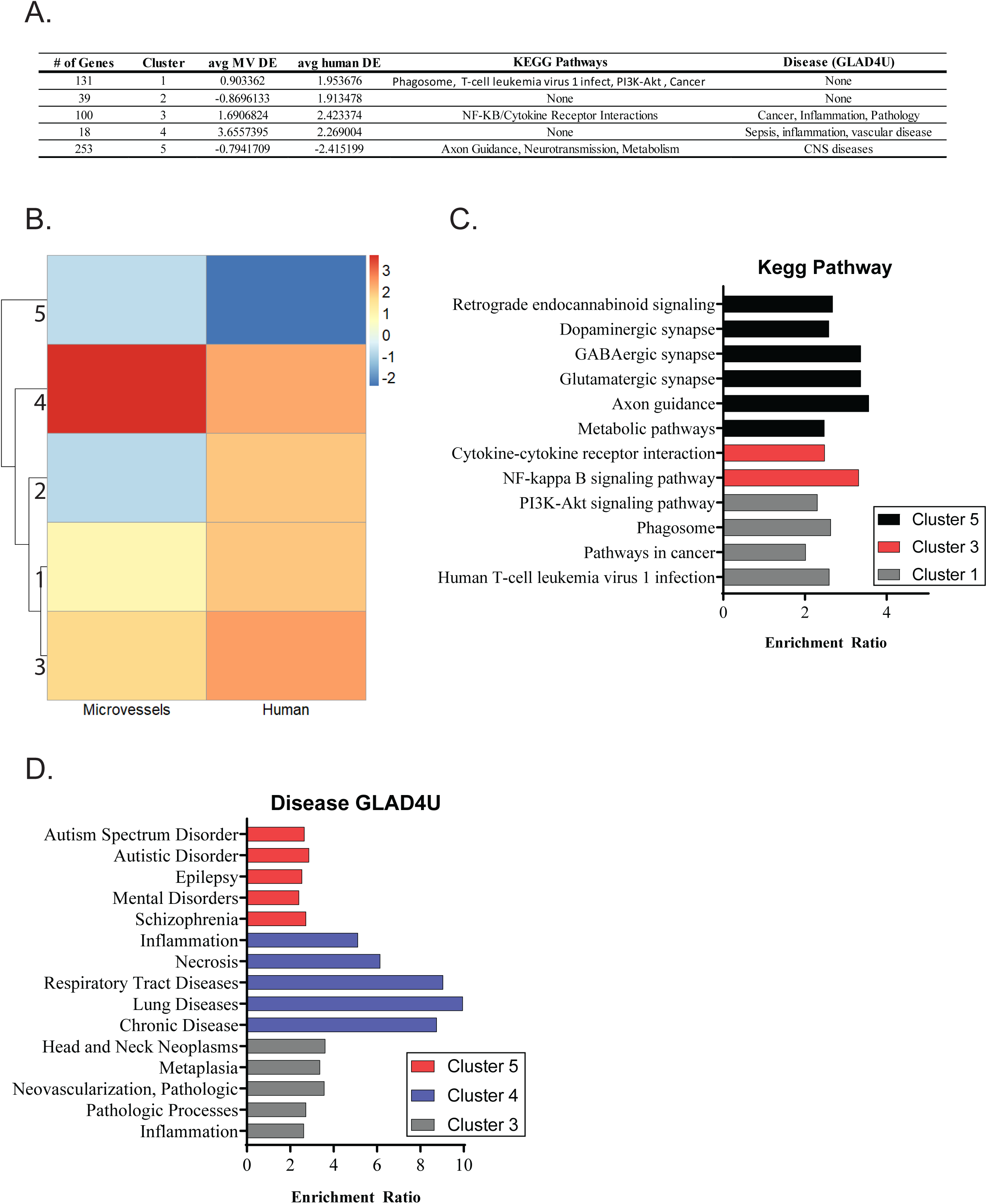
A) Table of clusters generated by k-means clustering with a seed set to 25. B) Heatmap of k-means clusters (k=5, seed = 25). C) GSEA analysis of genes in each cluster using a pathway database (David Kegg). Clusters 2 and 4 resulted in no significant pathway predictions. D) GSEA analysis of genes in each cluster using a human disease database (GLAD4U). Clusters 1 and 2 resulted in no significant disease predictions.

**Supplemental Figure 5:**
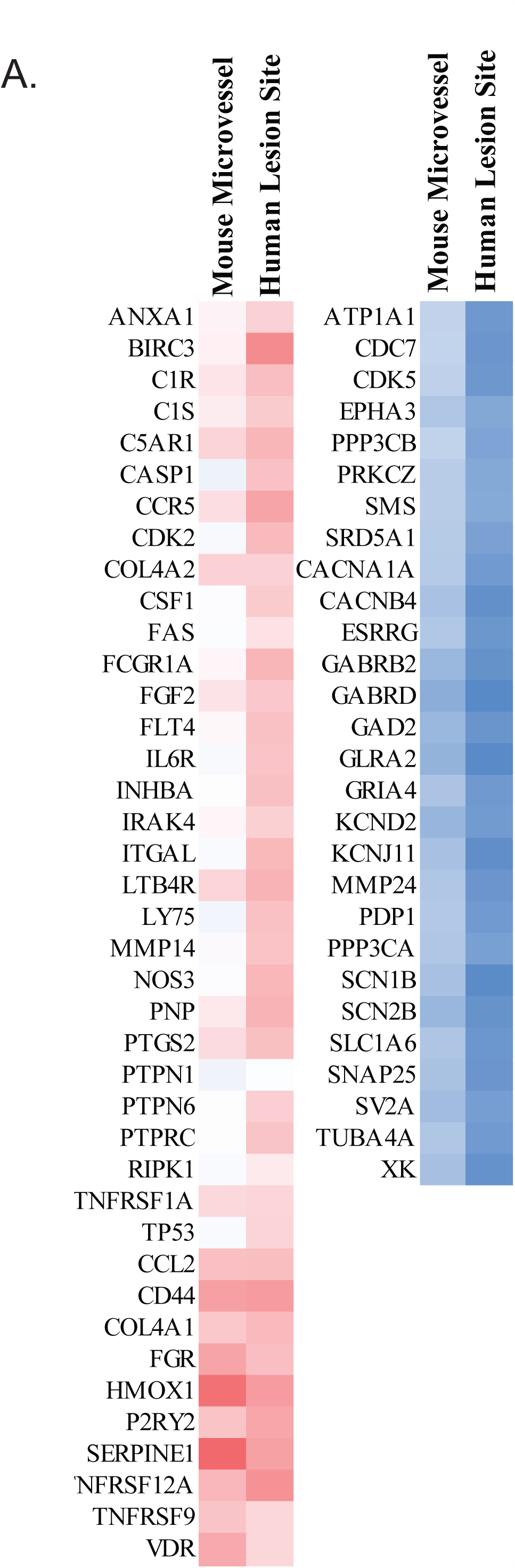
A) Heatmap of differential gene expression Z-scores of druggable gene targets in mouse microvessels and human lesion-sites identified through IPA.

**Supplemental Figure 6:**
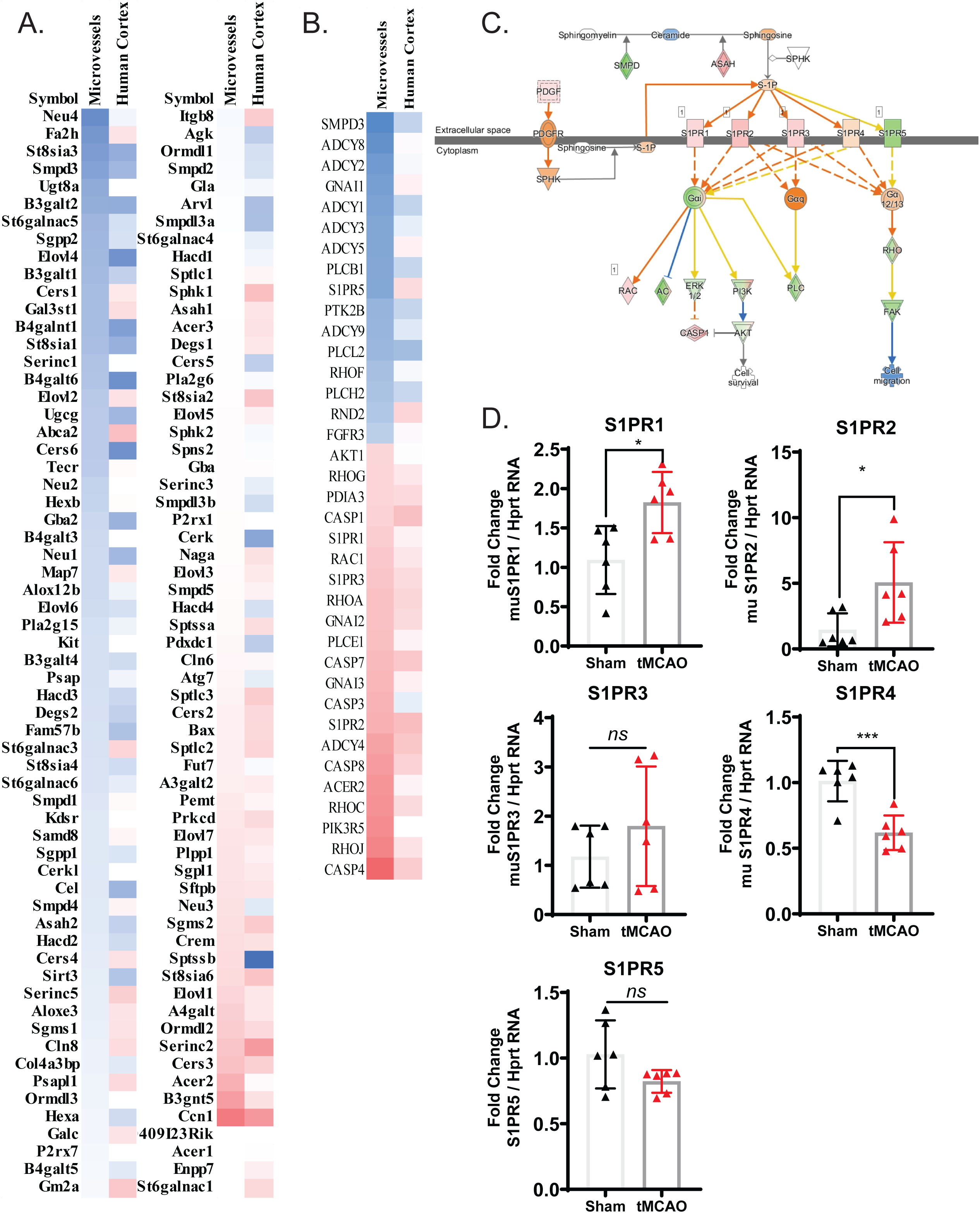
A) Heatmap of differential gene expression Z-scores for sphingolipid metabolic processes (GO: 0006665) including mouse microvessels and human lesion-sites. B) Heatmap of differential gene expression Z-scores of sphingosine receptor signaling components. C) Schematic representation of sphingosine receptor signaling generated by IPA and overlaid with differential gene expression in microvessels after tMCAO. D) qPCR validation of sphingosine receptor transcripts. (*p<0.05;***p<0.001; ns = not significant).

**Supplemental Table 1:** General characteristics of mouse microvessel and human lesion site datasets.

| Dataset | Cell/Tissue Type | Disease Model | Profiling Method | Lab Details | NIH Accession Number |
| --- | --- | --- | --- | --- | --- |
| Mouse Microvessels | Cortical microvessels | Transient MCAO | RNA-seq | Teresa Sanchez - Weill Cornell Medicine | GSEXXX |
| Human Lesion-site | Cortex (lesion-site) | Stroke Patients (post-mortem) | RNA-seq | Jonas Frisen - Karolinska Institutet | GSE56267 |

**Supplemental File 1:** Excel file containing clustering data from 541 shared genes between mouse microvessels after tMCAO and human lesion sites after stroke.

**Supplemental File 2:** Excel file containing correlations between mouse microvessels after tMCAO and human lesion sites after stroke and gene lists from IPA canonical pathway categories: Neuroinflammation, Protein Kinase A, and Dopa-DARPP32 signaling.

